# Machine Learning-aided Discovery of Novel Chemotype Antagonists for G Protein-coupled Receptors: The Case of the Adenosine A_2A_ Receptor

**DOI:** 10.1101/2023.03.31.535043

**Authors:** Jonas Goßen, Rui Pedro Ribeiro, Dirk Bier, Bernd Neumaier, Paolo Carloni, Alejandro Giorgetti, Giulia Rossetti

## Abstract

Identifying the correct chemotype of ligands targeting receptors (i.e., agonist or antagonist) is a challenge for *in silico* screening campaigns. Here we present an approach that identifies novel chemotype ligands by combining structural data with a random forest agonist/antagonist classifier and a signal-transduction kinetic model. As a test case, we apply this approach to identify novel antagonists of the human adenosine transmembrane receptor type 2A, an attractive target against Parkinson’s disease and cancer. The identified antagonists were tested here in a radioligand binding assay. Among those, we found a promising ligand whose chemotype differs significantly from all so-far reported antagonists, with a binding affinity of 310±23.4 nM. Thus, our protocol emerges as a powerful approach to identify promising ligand candidates with novel chemotypes while preserving antagonistic potential and affinity in the nanomolar range.

## Introduction

Human guanine nucleotide-binding protein-coupled receptors (hGPCRs) represent one of the five protein families commonly targeted by prescription drugs, together with ion channels, kinases, nuclear hormone receptors, and proteases^1^. New possibilities for hGPCR drug discovery have emerged exploiting the modulation via allosteric sites, as well as an improved understanding of receptor activation mechanisms. The former enables targeting protein regions, which are less conserved between family subtypes than the primary binding site, leading to significant specificities.^2^ The latter, knowledge about activation mechanisms, can be facilitated manifold in virtual screening to not only identify key residues that can be targeted to modulate receptor function^2,3^, but also to sample and fix receptor conformations to ‘bias’ the signaling pathways associated with that subset.^4,5^ Also, the knowledge of pharmacological cellular-pathway-dependent parameters such as cAMP accumulation, calcium flux, ERK phosphorylation, arrestin recruitment, and G-protein interaction is now recognized as key to link *in-vitro* bioactivities with *in-vivo* effects.^4,6^

These developments are further extended by Quantitative Systems Pharmacology (QSP) multi-scale models that combine computational and experimental methodologies.^7^ The drug concentration profile in the blood, the site of action, and cellular signaling downstream of the targeted sites are a few examples of the key quantitative and qualitative factors that can be integrated and predicted using such models.^8,9^ QSP can clarify, validate, and apply new pharmacological principles to the creation of therapies that relate the kinetics of drug-target interactions to the cellular response that follows in the context of the (diseased) physiological state.

Here, we address the question of whether it is feasible to combine QSP-like approaches and structural data to select ligands with novel chemotypes, specifically ligands not mimicking natural substrates, that however retain a high probability to activate or inactivate the GPCR, and a promising dose-curve response of the relevant pathway.

As a test case, we focus on the human Adenosine Receptor type 2A (A_2A_R), an important target for pharmacological intervention (see Figure 1A). A_2A_R is a member of the subclass of rhodopsin-like receptors (class A) of G-protein coupled receptors, for which many functional data (e. g. IC_50_, EC_50_, K_i_, K_D_) can be exploited. The ligands, classified as full/partial agonists, neutral antagonists, as well as inverse agonists, depending on its potential degree of receptor activation^3^, can act as pharmaceutical therapeutics targeting the A_2A_R.

**Figure 1:**
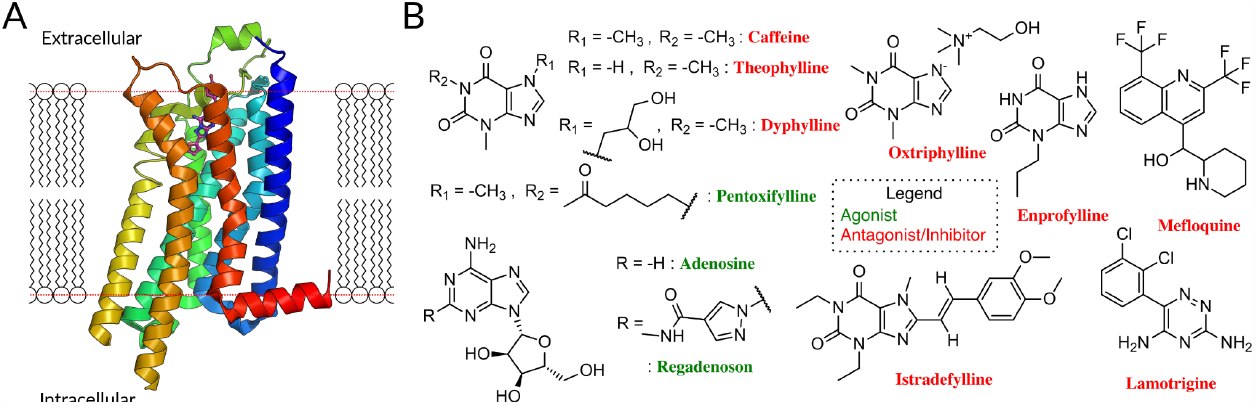
(A) The A_2A_R, like other GPCR, consists of seven transmembrane helices linked by three extra- and intracellular loops (ECL/ICL) and an amphipathic helical segment (membrane plane represented by the dotted red line)^16^. The latter lies parallel to the cytoplasmic membrane surface, connecting the membrane domain with its long C-terminal tail. (B) FDA-approved drugs for A_2A_, antagonists^10^ (red names) and agonists (green names) are highlighted. Antagonists are primarily derivatives of xanthine (caffeine, theophylline, dyphyline, oxtriphylline, enprofylline, pentoifylline, istradefylline), quinoline (mefloquine), triazine (lamotrigine). The xanthines theophylline, caffeine, and enprofylline are unfortunately rather unspecific binders, sharing affinity between A_2A_, A_1_ and partly A_3_.^17^ Oxtriphylline, the choline salt of theophylline, and dyphylline share this unspecificity to the A_2A_ and are mostly used as inhibitor for phosphodiesterase 4A.^18^ Pentoxifylline, also a theophylline derivate, shares activity between A_2A_ and A_1_ and targets phosphodiesterase enzymes^19^. Mefloquine is an antimalarial agent used in the prophylaxis and treatment of malaria^20^, which is only a moderately selective adenosine A_2A_R antagonist.^21^ Lamotrigine is an antiepileptic used to treat some types of epilepsy and bipolar disorder, while having estimated affinity for 17 different targets.^22^ Finally, despite suggested improvements^23^, istradefylline suffers from poor photostability.^23^

8 of 11 FDA-approved drugs targeting A_2A_R are antagonists, along with compounds undergoing clinical trials (see Figure 1B)^10^. They are used to treat different diseases, mostly symptoms of neurodegenerative diseases like Parkinson’s^11,12^, as well as cancer^13,14^. Although these molecules are chemically diverse (see Figure SI 1), they all (i) lack the sugar moiety and (ii) feature a mono-, bi-, or tricyclic structure, which mimics the adenine part of adenosine. They are classified as xanthines and non-xanthines.^15^

Unfortunately, the existing antagonistic FDA drugs have issues that limit their domain of applicability (see caption of Figure 1 for further info) and many efforts are made to identify new ligands. So far, structure-based virtual screening has managed to achieve some success in identifying antagonists specific for A_2A_ receptors: several of them are active in *in-vitro* and, in one case, in a rat model (see Table 1)^24–28^. However, three major challenges need to be faced: (i) the difficulty to identify antagonists specific for this class of the receptor and not for other adenosine receptor subclasses, such as the A_2B_ receptors (59% similarity)^29^. This is crucial to avoid affecting the function of untargeted subtypes, possibly causing side effects^30^. (ii) the challenges in efficiently predicting the activity of the antagonists. Although there is indeed an increasing amount of activity data, structure-activity model approaches are often hampered by noise due to undersampling: Molecules known to be active (or inactive) are outnumbered by the vast amount of possible chemical features that might determine binding.^31^ (iii) the difficulty in predicting their mode of action in cellular pathways. Adenosine signaling is indeed widespread in several cascades and ligands acting at these receptors exert a broad spectrum of physiological and pathophysiological functions. However, predicting the effects of A_2A_R antagonists on A_2A_R cascade is not usually part of virtual screening.^32^ Similar challenges may be encountered across other GPCRs’ other than A_2A_R.

**Table 1:**
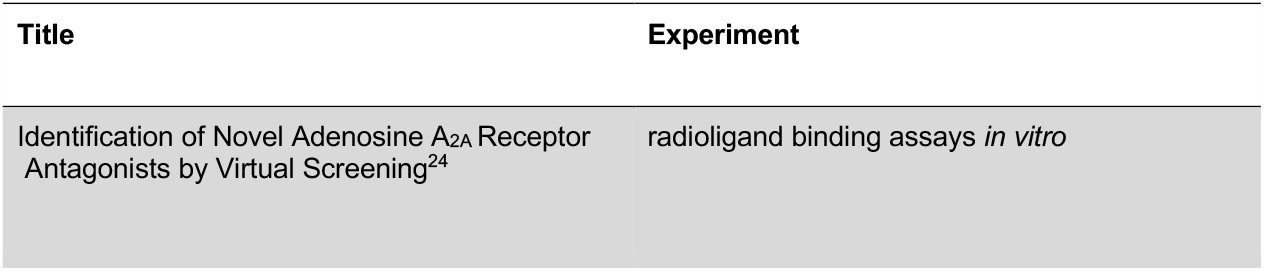
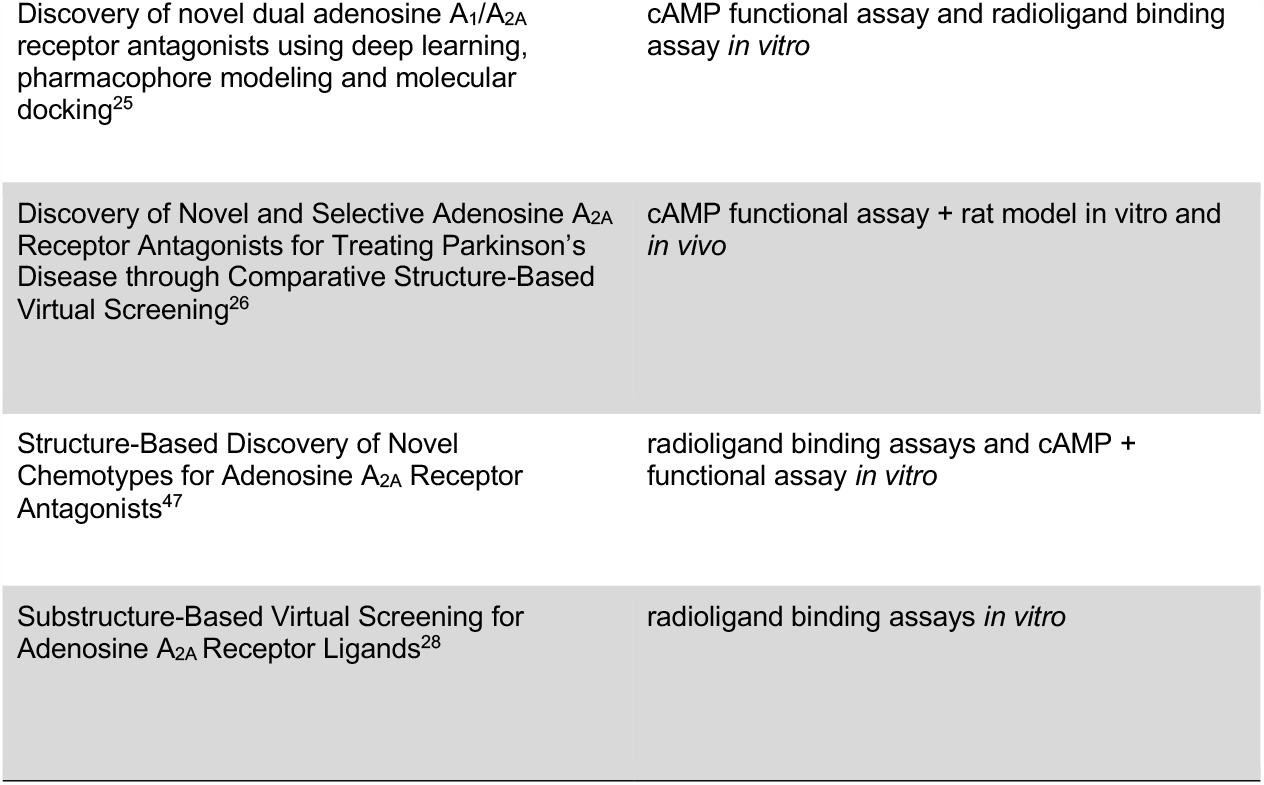
Comparison among Virtual Screening campaigns with added experiment type for A_2A_R reported in the literature.

Novel chemotype antagonists, tailored to the inactivated structure of the A_2A_R and not featuring common chemical aspects of general antagonists for adenosine receptors’ class, could be used in principle to solve the problem of specificity, i.e., point (i) above. However, with a novel chemotype, it is even more difficult in virtual screening campaigns to predict their activity or antagonist power (i.e., their ability to inhibit A_2A_R), through structure-activity type of modeling, as discussed in point (ii) above. Furthermore, such antagonist power needs in principle to be evaluated in a given cellular cascade, i.e., point (iii) above; This is usually beyond the domain of applicability of virtual screening campaigns.

To find novel chemotype antagonists we performed an unbiased virtual screening campaign targeting the inactivated structure of A_2A_R. To address the issue of predicting antagonistic power of novel chemotypes we used machine learning methods (ML), which are also robust to noisy data.^33^ And, to evaluate the impact of the selected antagonist candidates within a given subcellular pathway, we implemented an in-house QSP approach in the virtual screening pipeline. From one side, ML has indeed emerged as a very powerful tool in computer-aided drug design^34,35^. It is able to predict drug contrary effects^36^, identify new inhibitors, ^37^ and predict subclass specificity for ligands of A_1_ and A_2A_ receptors^38^, and even classify ligands’ mode of action for the β_2_ adrenergic receptor (AR).^39^ From the other side, QSP approaches like signal transduction models, comparably to ML, have also found extensive usage in the field. They are used to model *in vitro* cAMP response curves for the D_2_ Dopamine antagonist^40^, to optimize chimeric constructs that target EGF receptors in terms of selectivity and efficiency^41^, and used to characterize the bronchodilatory response of a 5-lipoxygenase inhibitor^42^. These successes encouraged us to develop random forest classifiers and combine it with our recently developed Structural Systems Biology tool^52^ within a virtual screening pipeline. The first allowed us to improve decision-making in selecting molecules with a higher potential to be antagonist, the second allowed us to predict the dose-response curves for the best-scored virtually screened antagonists, within the A_2A_R cascade. In this way, the final candidate selection was enormously improved, with respect to a canonical virtual screening campaign. By testing their binding affinity through in-vitro radio-ligand binding assays, we showed that one of our molecules is indeed an antagonist. The candidate shows a dose-response curve similar to the A_2A_ ‘gold standard’ inhibitor named ZM241385^53^, which is not only highly affine (K_D_=1.2 nM)^45^, but also selective for the A_2a_R subtype. Although the candidates’ affinity is sub-micromolar, its chemotype differs heavily from those reported in the ChEMBL database^46^ for the ligands targeting A_2A_ proving that QSP-like approaches, AI and structural data are a powerful combination to find novel chemotype molecules with proven antagonistic impact on the selected GPCR.

## Results

Data collection and characterization of known Agonists/Antagonists Compounds.

### Data Collection

Currently, (i.e., February 2023) there are 66 solved structures of the adenosine receptor type 2A (see Table SI 1).^48,49^ While 11 structures are bound to an agonist and represent an intermediate state in between inactive and active state, two structures are crystallized in a full active conformation. These full active conformations were achieved by binding the receptor to an engineered G-protein (mini-G_s_) in the case of the X-ray structure with ID 5G53^50^, and to an heterotrimeric G-protein (mini-G_s,_ βγ subunits and a nanobody Nb35) with the agonist NECA (see Figure 2A, left) in the case of cryo-EM solved structure 6GDG.^51^ Of the remaining structures, 53 are bound to an antagonist, of which 28 are the low nanomolar affinity ligand ZMA (see Figure 2B, left).

**Figure 2:**
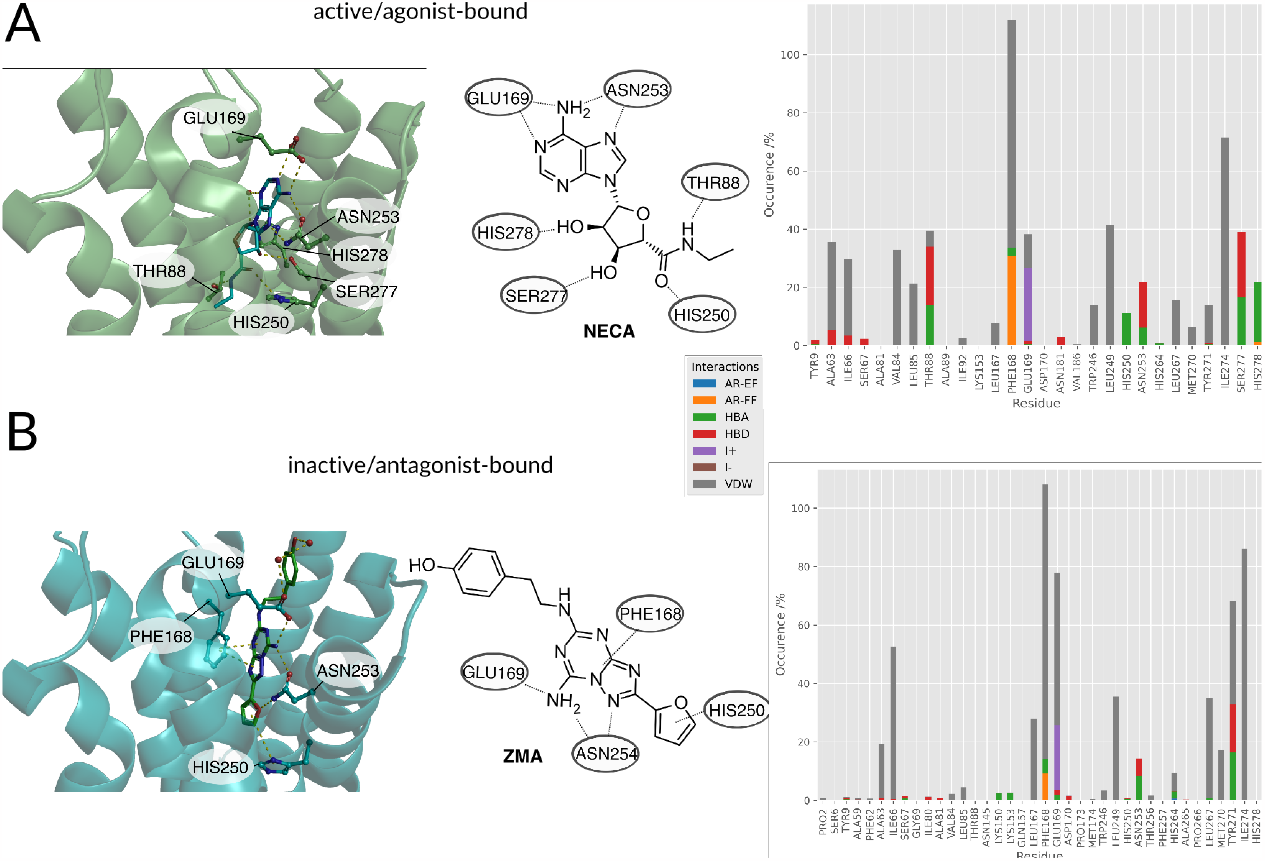
(A, left)Agonist/(B, left)Antagonist binding poses and docking histograms. Comparison between the binding conformations of an agonist NECA (PDBID:2YDO^57^) and antagonist ZMA (PDBID:4EIY^58^) of the A_2A_R in the left panels. Only polar interactions are highlighted in respective figures. (A, right/ B, right): PLIF of agonists and antagonists found in literature docked to the respective active and inactive receptors.

### Data characterization: Binding patterns differences of Agonists and Antagonists in Crystal Structures

Both agonists and antagonists make similar polar interactions between their adenine-like moieties with ASN253^6×55^ and GLU169^ECL2^ (see Figure 2A/B, left). The main difference is given by the fact that agonists contain a ribose group, which extends deeper than ZMA into the orthosteric binding pocket. There, the ribose moiety performs polar interactions with the HIS278^7×42^ and SER277^7×41^ residues in helix 7 and makes hydrophobic contacts with residues in helix 3 through THR88^3×36^. While all co-crystallized ligands engage helix 3 and helix 7 through these residues, some antagonists do also have interactions with them, but never at the same time.^52^ Lacking these interactions, the inverse agonist ZMA seems to be capable of preventing a crucial conformational change in helix 5 and, therefore, prevent the transition to the active state of the receptor.^53^ The ribose or the ribose-like moiety is part of the majority of known agonists (see Figure SI 2) and increases the affinity at all adenosine receptor subclasses. The substituted adenine moiety is mainly harnessed to determine the subtype selectivity for the generation of specific subtype agonists.^53,54^ Non-nucleosidic agonists are in the minority, but they represent an alternative to the generally long and complex synthetic route for substituted nucleosides.^55^ These consist mainly of purines or pyrimidines scaffolds and mimic the agonistic interactions of adenosine. In the case of A_2A_ agonist BAY-60-6583 a carboxamide function is mimicking the ribose-like interactions.^56^

Summarizing the crystallographic data, the binding patterns that leads to the inhibition or the activation of the receptor can be very diverse but some key interactions that distinguish agonist from antagonist can still be identified: the agonists engage with helix 3 (e.g., THR88^3×36^) and helix 7 (e.g., HIS278^7×42^, SER277^7×41^). Antagonists are less restricted in their explicit interaction profile as long as they block agonist access to the residues mentioned above.

### Data Enrichment

Next, we added additional molecules to the ensemble of approved drugs and co-crystallized molecules by including a library of known agonist/antagonist of the A_2A_. These were collected from GuideToPharmacology^59^, GPCR database^49,60,61^, ChEMBL ^46^, and Drugbank^10^ (see Figure SI 3, left). Lacking experimental data for these small molecules, we estimated the complex conformation via molecular docking (see Methods). Distinct differences between the binding patterns of known agonists and antagonists emerged in a similar fashion to the crystal conformations (see Figure 2, right): Hydrogen bond donor interactions of SER277^7×41^ and HIS278^7×42^ are very specific to the agonists poses. The hydrogen bond donor interaction with ASN253^6×55^ is also far more frequent in agonists’ binding poses. While the ionic interaction with GLU169^ECL2^ can also be found in around 50% of antagonist’s poses and in more than 80% of the agonists’ poses. Another very distinct feature is the agonist’s hydrogen bond donor interaction with THR88^3×36^, which is not as frequent in the antagonists’ poses. This is due to the deep placement in the binding pocket and can therefore only be reached by mostly linear antagonists. These interactions are similar to the ones found in the crystals structures already published for the adenosine receptor and mentioned above. Main anchoring interactions are the salt bridges formed with GLU169^ECL2^ and π-π-stacking (aromatic face to face) interactions with PHE168^ECL2^ in both active and inactive structures. The biggest differences can be observed in the contacts with SER277^7×41^ and HIS278^7×42^, which act as hydrogen bond acceptors for the ribose-moiety of the agonists. Differences between the crystal structure and the docking poses can be explained by the limitations of the docking approach: the protein was kept in one rigid state allowing flexibility only for the small molecule binder. Crystallographic effects, like contacts with the crystal image in the unit cell are also a possible source of different behaviors on passing from the X-ray to the docked structure. After confirming differences between agonist and antagonist docking poses, which are similar to the experimental conformations, the docking poses and virtual library of agonists/antagonists was used to build two random forest classification models.

### Fingerprints and Random Forest Classifiers

Having found distinct differences in the docking poses of Agonists compared to Antagonists, we can now make use of this finding with a classical machine learning approach: Two random forest models were built for the distinction of agonists from antagonists. One was based on the chemical graph representation (extended-circular fingerprints, ECFP4 hereafter) of all the known adenosine receptor ligands available to us (ECFP4 Based Random Forest). The second one was based upon the interaction fingerprint of the same library analyzed above, docked to the receptor in an active, inactive and intermediate state of activation (PLIF Based Random Forest). The five-fold cross-validation performance evaluation metrics are listed for both models in Table SI 2 and Figure SI 4. The baseline performance of both the classifiers is very high (>.9 accuracy). Due to the high baseline-performance, hyperparameter optimization was not applied to avoid overfitting effects.^62^ To evaluate which ligand features (ECFP4 based) and interactions (PLIF based) are responsible for the decision we analyzed the models by calculating the SHAP (**SH**apley **A**dditive ex**P**lanations)^63,64^ value impacts on the model output (see Methods for further details) of the available data in the following paragraphs.

#### Features evaluation

The SHAP value of the eight most important features for the ECFP4 based classifier are plotted in Figure 3A. Features #190 and #454 are encoding for fragments centered on an ether function and a carbon bound to a hydroxyl group, respectively (see Figure 3B). Both features are very reminiscent of the ribose moiety bound to nucleobase adenine (as found in most agonists, see Figure SI 2). Features #1257 and #3535 both show fragments of a hydroxyl function needed for hydrogen bond interactions. Features #3314 and #3709 show nitrogens as part of an aromatic ring system. These features (including feature #2020, which shows a sp^2^ hybridized carbon) can also be found in parts of the nucleobase adenine. Lastly feature #3067 shows sp^3^ hybridized carbon, which seems rather unspecific, but is present in all agonists.

**Figure 3:**
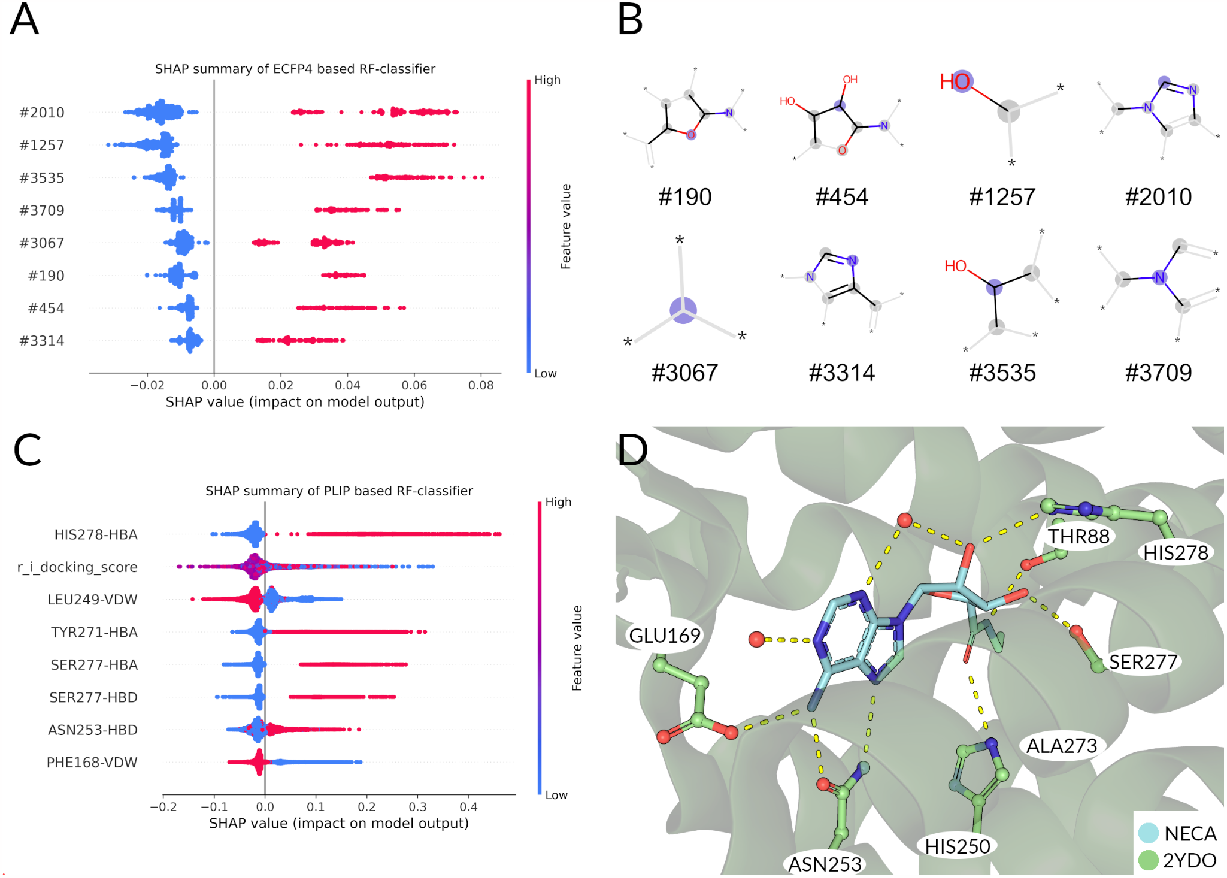
(A) Feature importance by SHAP value calculation on training set. Each point corresponds to the impact of the morgan fingerprint feature to the classification as ‘agonist’. Furthermore, a ‘high’ value (colored red) corresponds to the presence, a ‘low’ value (colored blue) to an absence of the corresponding feature in the plot. For all the features shown, a presence of a feature contributes to the classification as agonist, while an absence indicates an antagonist. (B) Visualization of bit fragments for fingerprint bits for classification by final ECFP4 RF model. (C) Feature importance by SHAP value calculation of PLIF based rf model shows most important features for agonistic classification. (D) Binding pose of NECA shown for reference of interactions. Water molecules are shown as red spheres.

Following the same procedure as in the ECFP4 based model, the features importance was analyzed using SHAP values (see Figure 3C). The most important features for agonist predictions are hydrogen bond interaction with HIS278^7×42^, TYR271^7×35^ and SER277^7×41^. These interactions are frequently found in the known A_2A_ agonists and mediated mostly by the ribose-like-moiety (see Figure 1A). Interestingly, the docking score is the second important feature for discriminating agonists from antagonists in this classifier (see Figure 3C and Figure SI 5). Unsurprisingly, the average agonist shows a lower glide score in the active state receptor compared to the inactive state (see Figure SI 5AB). The ligands with the lowest docking scores show to be primarily agonists. This observation is mirrored in the SHAP values for the docking score: A low value predicts mostly agonists, while a mediocre value is more common in the antagonist docking poses. Van-der-Waals interactions with PHE168^ECL2^ is most indicative of the antagonists’ poses, as this interaction (paired with an aromatic face-face interaction) is often used to ‘anchor’ the aromatic core of most antagonists in the binding pocket. Also, the presence of the LEU249^6×51^-Van-der-Waals interaction predicts antagonists. The anchoring interaction performed by ASN253^6×55^ is also more likely to occur in agonists’ (see Figure 3D) poses compared to the antagonists’. Summarizing, in the ECFP4-based RF model the existence of ribose-(e. g. #190, #454) and adenine-like (e.g., #3314 and #3709) fragments are among the top relevant features for the decision of the model. Instead, the PLIF-based RF model’s features are more based upon interactions that are crucial for the activation/inhibition of the receptor (e. g. HIS278^7×42^, TYR271^7×35^ and SER277^7×41^), which are not linked intrinsically to a particular chemical structure. Therefore, since our scope is to find novel scaffolds with antagonistic power, we prioritized the PLIF-based classifier.

### Virtual screening and filtering: Selection of Molecules for the Experimental Testing

After analysis of the random forest models, the ‘Schrodinger’s Virtual Screening Workflow’ (see methods for details) was used for the screening of 403716 structures against the crystal structure of A_2A_ (PDB_ID: 4EIY^58^). By docking to the receptor in the inactive state, the probability of finding an antagonist is increased.^65^

#### Filter 1: RF classifiers

307 molecules representing the top 1% of scoring ligands resulting from the virtual screening workflow were obtained and filtered with the RF classifiers. Figure SI 6 shows histograms of the estimated antagonist power probability. All of the top 1% molecules (307) selected by virtual screening show a high average probability to be antagonists by the PLIF-based RF classifier. Therefore, we chose to neglect molecules with a probability lower than 0.84.

#### Filter 2: Predicting Dose-Response Curves using Systems Biology Simulations

We have used an in-house developed Structural Systems Biology (SSB) approach^43^ to simulate the signaling pathway (or cascade) of the Adenosine 2A receptor to reproduce, in-silico, the subcellular dynamics of cAMP upon ligand-target binding. The aim was to obtain the predicted dose-response curves from the SB simulations to rank qualitatively the potency of the twelve most promising molecules obtained from the virtual screening. See the paper from some of us for detailed protocols.^43^ We made sure the selected compounds showed a pK_d_ lower than 5.5 (corresponding to 10 *μ*M) in the KDeep predictor^66^.

#### Clustering

We clustered the molecules according to their ECFP4 fingerprint to find a most diverse subset of molecules (see methods). Specifically, the extended circular fingerprints were generated and the pairwise Tanimoto similarity between the molecules was used to construct a distance matrix (see Figure SI 7 for a TSNE visualization). The latter was used for hierarchical clustering and we chose 90 clusters based on silhouette score^67^.

Of the different clusters the final molecules were chosen by visual inspection, comparing the docking pose to the crystal ligand ZMA and showing favorable interactions with the receptor. In Figure 4A the final selection of molecules’ 2D structures is shown with the corresponding dose-response curves; Although all the molecules show predicted behaviors in the high micro to low-nanomolar range (see Figure 4B), we can see that there is a distinct difference between the first (JG_03) and last compound (JG_10).

**Figure 4:**
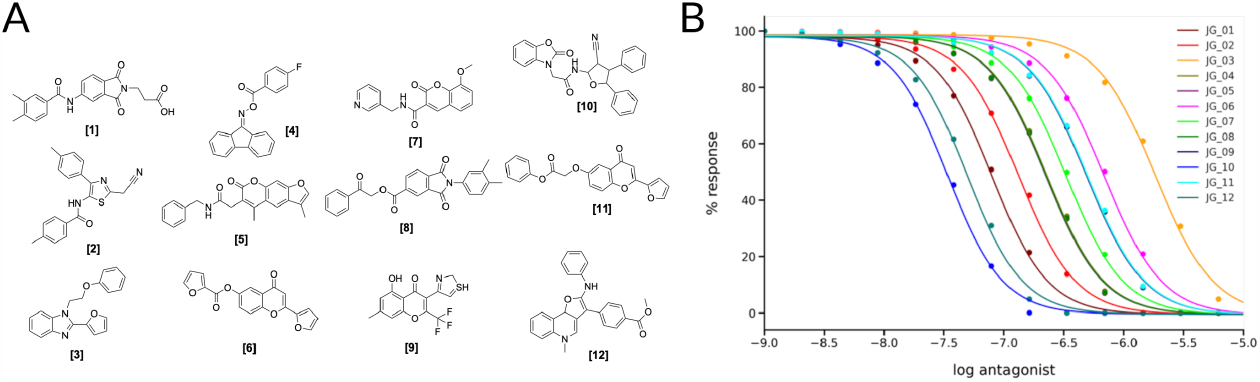
(A) 2D Structures of final selected molecules by the virtual screening process. (B) ligand-target dose-response curves of the 12 molecules obtained through the SSB toolkit by using the pK_D_ values to initiate the SB simulations. The response is obtained by calculating the change in the concentration of cAMP in function of simulation time and ligand concentration. For the simulations we used a receptor concentration of 2 μM, a range of ligand concentration between 10^−3^ μM and 10^3^ μM, and an integral time step of 1000. The affinity estimated by KDEEP^66^ of the compounds are listed in Table SI 4.

### Binding Experiments and Induced Fit docking

The binding affinity of the compounds visible in Figure 4A were tested in competition experiments against the radio ligand ZMA (see Figure 5A and Figure SI 8). And indeed, the most promising molecule found is JG-10, which shows a binding affinity of 310 ± 23.4 nM and a similar dose-response curve as predicted (see Figure 5B). Additional Induced-Fit docking was performed on the 12 tested compounds, to allow for sidechain flexibility of the poses. The induced fit optimized scores are listed in Table SI 5 and the docking pose of compound JG-10 is shown in Figure 5CD. There are four polar interactions between the receptor and the ligand: Two hydrogen bonds are formed with the hydroxyl group of the dihydrobenzooxazol moiety. Both are backbone interactions with the peptidic nitrogens of PHE168^ECL2^ and GLU169^ECL2^ at distances of 2.8 and 2.2 Å respectively. Additionally, PHE168^ECL2^ forms a π-π interaction with both ring systems of the ligand with PHE168^ECL2^ at a distance of 3.57 Å. These interactions are with the extracellular loop 2, which is specific for the A_2A_ and is found to contribute to subtype specificity in other ligands. The furan moiety bound nitril function forms a hydrogen bond with the side chain N of ASN253^6×55^ at a distance of 1.96 Å. The dihydrobenzooxazol moiety also forms hydrophobic contacts with ILE66^2×63^, TYR271^7×35^ and LEU267^7×31^ (see Table SI 6). Additional hydrophobic contacts are formed by the two phenyl moieties bound to the furan core, which form interactions with VAL84^3×32^, LEU85^3×33^, ALA63^2×60^, and LEU249^6×51^ (see Table SI 6).

**Figure 5:**
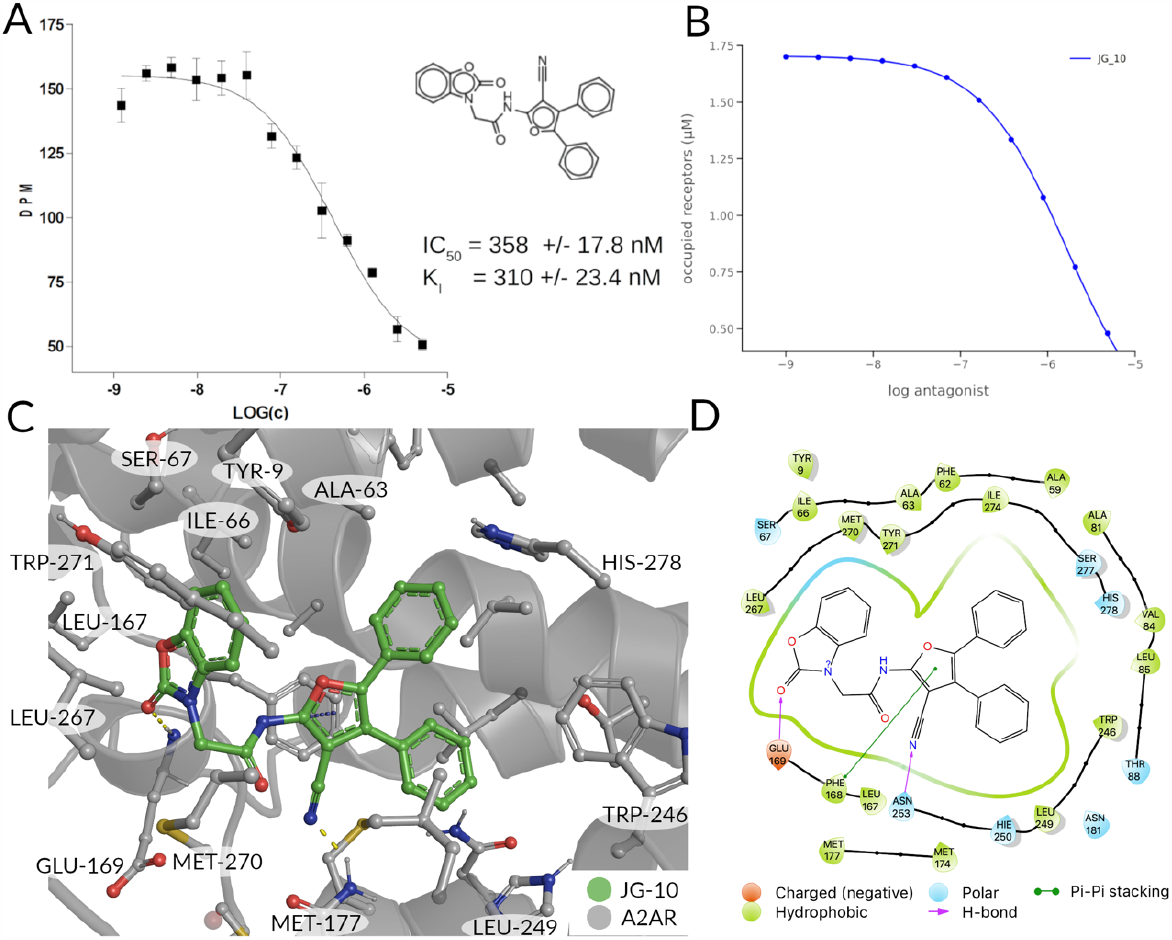
(A) Radioligand-binding displacement assay of novel binder (JG-10). (B) ligand-target binding curve of JG_10 obtained by using the pKd value to calculate the fraction of occupied receptors at equilibrium. (C,D) Induced-Fit docking pose of JG-010 in A_2A_R (4eiy) in 3/2D. Polar interactions are shown in yellow (H-Bonds) and orange (π-π stacking interactions).

### Checking for the novelty of top selection

The tanimoto similarity metric between the extended circular fingerprints with radius 2 (ECFP4^68^) fingerprints of known binders of the adenosine receptor deposited in the ChEMBL database^46^ and the one used in the experimental testing were calculated using a python script (see Table SI 7). Compared to their most similar structure found in ChEMBL database these molecules show some significant differences in their chemical structure. Most promising scaffold JG-10 differs greatly from its most similar molecule found in ChEMBL with a T_c_ below 0.4 and can therefore be denoted as a ‘novel’ compound.

### Conclusion

We have presented a novel virtual screening protocol that integrates a ML model able to classify candidate receptor binders based on their antagonist/agonist potential. In this work, we have focused on the A_2A_R. Specifically, two random forest models were built by learning from antagonists and agonists found in literature to filter out agonistic binding patterns. One was based on the chemical graph representation (extended-circular fingerprints, ECFP4) of all the known adenosine receptor ligands available in publicly available databases. The second one on an interaction fingerprint of the same library docked to the receptor in an active, inactive and intermediate state of activation. While both models accurately classify the available data, they differ in their underlying representations. The ECFP4-based model is built on well-established chemical space. In contrast, the PLIF-based model captures subtle features of protein-ligand interactions that chemical graph-based approaches may only partially represent. We chose to use the PLIF model to explore novel chemical space and potentially discover new ligands that are not limited by the explicit chemical structure in the training set. The RF classifier predicts the ligands to be antagonists with a high probability. Next, the dose-response curves of the most promising compounds were predicted using an in-house developed Structural Systems Biology (SSB) approach, further guiding the compound selection. The final selection found by the virtual screening pipeline applied in this work was validated with an in-vitro radio ligand binding assay experiments by our collaborators. One of the compounds JG-10 has shown an affinity (K_I_) of 310±23.4 nM. The most promising compound has a distinct chemotype compared to the most similar molecule of the ChEMBL database (T_c_ (ECFP4) = 0.33). The procedure described here could be applied straightforwardly to other GPCRs and even other types of membrane receptors.

## Materials and Methods

### Data Preparation and Fingerprint Generation

The interaction fingerprints were generated with an in-house-python script utilizing the interaction fingerprint implementation of the ODDT module^68^. The interaction fingerprint captures: (i) whether hydrophobic residues are in contact with the ligand; (ii) whether aromatic residues are oriented face to face; (iii) whether aromatic residues are oriented edge to face; (iv) whether a residue provides hydrogen bond acceptor(s); (v) whether a residue provides hydrogen bond donor(s); (vi) whether the residue provides hydrogen bond acceptor(s); (vi) whether a residue provides a salt bridge (protein positvely charged); (vii) whether a residue provides a salt bridge (protein negatively charged); (viii) whether a residue provides a salt bridge (ionic bond with metal-ion). Additionally, to the 8 bits of the PLIF for each molecule, the glide SP docking score was input for the algorithm.

### Random Forest Classifier Training

We used Schrödinger’s ligprep tool to find the most probable protonation state for the gathered molecules from A_2A_ specific binders from GuideToPharmacology^59^, GPCR database^49,60,61^, ChEMBL ^46^, and Drugbank^10^) molecules at pH 7. For the ECFP-based classifier we translated

For the PLIF based random forest classifier, we considered the best-scored five binding poses of each molecule docked to the three receptors (4EIY^58^, 5G53^50^ and 2YDO^53^) species to address the challenge of the docking approach to select the most physiological-like conformation.^69^

The models’ performance was evaluated by a cross-validation strategy. A 5-fold cross validation strategy was applied, in which scikit-learn’s StratifiedGroupKFold method was used to ensure no molecule pose was present in the training or the test splits. Model performance was evaluated by calculating the recall, precision, and f1 metric, which are defined below as implemented in the scikit-learn python package. While precision is the ability of the classifier not to label as positive a sample that is negative, recall is the ability of the classifier to find all the positive samples. The F1 can be understood as the harmonic mean between precision and recall and evaluates overall performance. The area under the receiver operating characteristic (ROC) curve is calculated by sklearn’ ROC AUC score function. The information about the plotted curve is therefore condensed into one value by calculating the area under the roc curve.

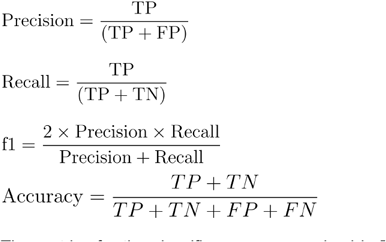

The metrics for the classifiers are summarized in Table SI 2. Insights into global model structure is gained by combining many local explanations of each prediction (SHAP python package). The SHAP package was used in a script using the final trained models.

### Virtual Screening and Induced Fit Docking

For the docking calculations we used a single crystal structure of the chimeric protein of A_2A_R-BRIL in complex with antagonist ZM241385 with a low available resolution and high overall quality (1.8 Å, PDB_ID: 4EIY^58^). We deleted intracellular loop 3 with apocytochrome b(562)RIL (Residues 1001-1106), as it was added to improve crystallization. Additionally, all waters beyond 5 Å distance to all HET groups denoted in the PDB-file and all Ligands except for ZM241385 were deleted. Hydrogens were added and bond orders were assigned using the Chemical Component Dictionary (CCD) database^70^. Schrödingers “Protein Preparation Wizard’’ was then used to generate the tautomeric states of all protein HET groups at a pH of 7±2. We added missing side chain atoms and optimized their conformations to the residues GLN148^ECL2^, GLU161^ECL2^, ARG220^6×22^ and ARG293^8×48^ with Schrödinger’s Prime program^71^. After optimization of the hydrogen bond orientations, all waters with less than 3 H-bonds to non-waters were deleted and the structure was minimized with the OPLS3 force field^72^. Convergence was assumed to be reached after the RMSD was equal or smaller as 0.30. We used “Schrödingers Virtual Screening Workflow” for the screening, which includes the following steps: (i) Initial Docking and ranking with the HTVS scoring function (ii) dock top 10 % (highest ranking) for docking with “Single Precision” (SP) Scoring(iii) dock top 10 % (highest ranking) for docking with ‘Extra Precision’ Scoring (XP). The remaining 307 top scoring ligands were then clustered to find a most diverse subset of molecules.

Induced Fit docking was performed using Schrödinger’s Maestro 2019.4.^73^ The centroid of the XP poses were used for the induced-fit calculation with a 10 Å inner box size (the automatic box size method was taken for the external box). Initial glide docking for each ligand was carried out on SP precision and side chains were trimmed automatically based on the B-factor, with the default 5 Å distance cutoff. B-factor cutoff was set to 40.0 and the maximum number of trimmed residues was set to 3. Prime refinement was carried out within 5 Å of docked poses with the optimize side chain option flag turned on. Glide XP re-docking into structures within 30.0 kcal/mol of the best structure, and within the top 20 structures overall was then carried out. The resulting ligand-complex orientations were evaluated by the IFD and Glide XP score.

### Clustering and TSNE

For the clustering of the final 307 molecules, we generated the ECFP4 (morgan fingerprints^74,75^, with a radius of 2 with a length of 4096 bits). We then generated the tanimoto distance matrix, for which we applied principal component analysis^76,77^(N=50) as implemented in scikit-learn. For visualization purposes we further reduced the data dimensions using T-distributed Stochastic Neighbor Embedding (TSNE)^78^. The dimensions of the distance matrix were reduced to 50 by PCA. For the final 2-dimensional embedding of the 307 virtual screening molecules fingerprints we chose a perplexity of 30, a learning rate of 200 and 5000 iterations.

### Experimental Details

The cells were grown adherently and kept in Ham’s F12 Nutrient Mixture, containing 10% fetal bovine serum, penicillin (100 U/mL), streptomycin (100 μg/mL), L-glutamine (2 mM) and Geneticin (G418, 0.2 mg/mL) at 37°C in 5% CO_2_/95% air. Cells were split two or three times weekly at a ratio between 1:5 and 1:20. For binding assays the culture medium was removed, cells were washed with PBS buffer (pH 7.4), scraped off and suspended in 1 ml PBS per dish, frozen in liquid nitrogen at a protein concentration of 6 mg/mL and stored at -80°C. Protein estimation used a naphthol blue black photometric assay^79^ after solubilization in 15% NH_4_OH containing 2% SDS (w/v); human serum albumin served as a standard.

Membrane preparation^80^: for radioligand binding experiments the frozen cell suspension was thawed and homogenized on ice (Ultra-Turrax, 30s at full speed). The homogenate was centrifuged for 10 min (4°C) at 600 g. The supernatant was then centrifuged for 60 min at 50.000 G, the membrane pellet was resuspended in 50 mM Tris/HCl buffer (pH 7.4) and stored at -80°C.

Binding experiments used membranes from CHO K1 cells stably expressing the human A_2A_R. Dissociation constant of [^3^H]ZM 241385 and the inhibition constant of not titrated ZMA, were obtained using [^3^H]ZMA (0.8 nM in competition experiments) as radioligand. Membrane homogenates with a protein content of 15 μg immobilized in a gel matrix (this method produces the same results as conventional separation techniques and will be published in detail elsewhere) were incubated with the radioligands in a total volume of 1500 μL 50 mM Tris/HCl buffer pH 7.4. After an incubation time of 70 minutes the immobilized membrane homogenates were washed with water and transferred into a scintillation cocktail (5 mL each, Ultima Gold, Perkin Elmer). The radioactivity of the samples (bound radioactivity) was measured with a liquid scintillation counter (Beckman, USA).

All binding data were calculated by non-linear curve fitting with a computer aided curve fitting program (Prism version 4.0, GraphPad Software, Inc., La Jolla, USA).

### Limitations

Although our SB approach is able to simulate dose-response curves and predict qualitatively the most potent ligand comparable with experimental data, one should keep in mind that the model was developed based on a previous model tuned in order to mimic specific experimental data for other GPCR.^81^ Here, future improvements of our model should involve fitting of kinetic constants to experimental data obtained for the A_2A_ receptor signaling pathway. Moreover, one should also take in mind that not only we used the affinity values as an exclusive parameter to calculate the fraction of activated receptors, but also that docking scores and experimental binding affinities are still poorly correlated.

Even though the affinity of the ligand to the A_2A_ is confirmed in experiment, there are several additional experiments needed to confirm the ligand selectivity by measuring the affinity towards other subtypes of the adenosine receptor such as the A_2B_ receptor. Also, the mode of action of the ligands predicted by our model was not explicitly measured in a functional assay.

Nevertheless, the implementation of prediction of dose-response curves using structure systems biology approaches within virtual screening campaigns, might support, in future, rational drug design projects.

## Acknowledgments

GR acknowledges the European Union’s Horizon 2020 Framework Programme for Research and Innovation under the Specific Grant Agreement No. 945539 (Human Brain Project SGA3), as well as the joint Lab “Supercomputing and Modeling for the Human Brain” of the Helmholtz Association, Germany and the European Union’s Horizon 2020 MSCA Program under grant agreement 956314 [ALLODD].

