## Supporting Information for "Machine Learning-aided Discovery of Novel Chemotype Antagonists for G Protein-coupled Receptors: The Case of the Adenosine A_2A_ Receptor"

*Table SI 1: Structural Data by Experiment.*

| Method | PDB | Res. (Å) | State | Degree active (%) | Name | Function | PDB Date | Ref. |
|---|---|---|---|---|---|---|---|---|
| X-ray | **8CU6** | 2.8 | Inactive | 1 | LJ-4517 | Antagonist | 31/08/2022 | [82] |
| X-ray | 8CU7 | 2.1 | Inactive | 1 | LJ-4517 | Antagonist | 31/08/2022 | [82] |
| X-ray | **8DU3** | 2.5 | Inactive | 1 | 21a | Antagonist | 10/08/2022 | [83] |
| cryo-EM | 7T32 | 3.4 | Inactive | 1 | ZM-241385 | Antagonist | 10/08/2022 | [84] |
| X-ray | **7EZC** | 3.8 | Active | 73 | UK-432,097 | Agonist | 13/04/2022 | [85] |
| X-ray | 7PYR | 2.6 | Inactive | 1 | Preladenant conjugate PSB-2115 | Antagonist | 02/03/2022 | [86] |
| X-ray | **7PX4** | 2.3 | Inactive | 1 | Preladenant conjugate PSB-2113 | Antagonist | 02/03/2022 | [86] |
| MicroED | 7RM5 | 2.8 | Inactive | 1 | ZM-241385 | Antagonist | 08/09/2021 | [87] |
| X-ray | **7ARO** | 3.1 | Inactive | 1 | CHEMBL124 345 | Agonist (partial) | 07/04/2021 | [88] |
| X-ray | 6LPK | 1.8 | Inactive | 1 | ZM-241385 | Antagonist | 25/11/2020 | [89] |
| X-ray | **6LPL** | 2.0 | Inactive | 1 | ZM-241385 | Antagonist | 25/11/2020 | [89] |

| Method | PDB | Res. (Å) | State | Degree active (%) | Name | Function | PDB Date | Ref. |
|---|---|---|---|---|---|---|---|---|
| X-ray | **8CU6** | 2.8 | Inactive | 1 | LJ-4517 | Antagonist | 31/08/2022 | [82] |
| X-ray | 8CU7 | 2.1 | Inactive | 1 | LJ-4517 | Antagonist | 31/08/2022 | [82] |
| X-ray | **8DU3** | 2.5 | Inactive | 1 | 21a | Antagonist | 10/08/2022 | [83] |
| X-ray | 6LPJ | 1.8 | Inactive | 1 | ZM-241385 | Antagonist | 25/11/2020 | [89] |
| X-ray | **6WQA** | 2.0 | Inactive | 1 | ZM-241385 | Antagonist | 18/11/2020 | [90] |
| X-ray | 6ZDV | 2.1 | Inactive | 1 | 2-Methyl-3-(4-methylthiazol-2-yl)-4-oxo-6-propyl-4H-chromen-7-yl acetate | Antagonist | 16/09/2020 | [91] |
| X-ray | **6ZDR** | 1.9 | Inactive | 1 | CHEMBL2030687 | Antagonist | 16/09/2020 | [91] |
| X-ray | 6S0Q | 2.7 | Inactive | 1 | ZM-241385 | Antagonist | 15/07/2020 | [92] |
| X-ray | **6S0L** | 2.7 | Inactive | 1 | ZM-241385 | Antagonist | 15/07/2020 | [92] |
| X-ray | 6PS7 | 1.9 | Inactive | 1 | ZM-241385 | Antagonist | 13/11/2019 | [93] |
| X-ray | **6JZH** | 2.3 | Inactive | 1 | ZM-241385 | Antagonist | 30/10/2019 | [94] |
| X-ray | 6GT3 | 2.0 | Inactive | 1 | imaradenant | Antagonist | 26/06/2019 | [13] |
| X-ray | **6MH8** | 4.2 | Inactive | 1 | ZM-241385 | Antagonist | 24/04/2019 | [95] |
| cryo-EM | 6GDG | 4.1 | Active | 100 | NECA | Agonist | 16/05/2018 | [51] |

| Method | PDB | Res. (Å) | State | Degree active (%) | Name | Function | PDB Date | Ref. |
|---|---|---|---|---|---|---|---|---|
| X-ray | **8CU6** | 2.8 | Inactive | 1 | LJ-4517 | Antagonist | 31/08/2022 | 82 |
| X-ray | 8CU7 | 2.1 | Inactive | 1 | LJ-4517 | Antagonist | 31/08/2022 | 82 |
| X-ray | **8DU3** | 2.5 | Inactive | 1 | 21a | Antagonist | 10/08/2022 | 83 |
| X-ray | **5WF6** | 2.9 | Active | 73 | UK-432,097 | Agonist | 21/02/2018 | 96 |
| X-ray | 5WF5 | 2.6 | Active | 73 | UK-432,097 | Agonist | 21/02/2018 | 96 |
| X-ray | **5OLV** | 2.0 | Inactive | 1 | CHEMBL167 1936 | Antagonist | 17/01/2018 | 97 |
| X-ray | 5OLO | 3.1 | Inactive | 1 | tozadenant | Antagonist | 17/01/2018 | 97 |
| X-ray | **5OM4** | 2.0 | Inactive | 1 | CHEMBL202 4114 | Antagonist | 17/01/2018 | 97 |
| X-ray | 5OLG | 1.9 | Inactive | 1 | ZM-241385 | Antagonist | 17/01/2018 | 97 |
| X-ray | **5OLZ** | 1.9 | Inactive | 1 | CHEMBL202 4114 | Antagonist | 17/01/2018 | 97 |
| X-ray | 5OLH | 2.6 | Inactive | 1 | vipadenant | Antagonist | 17/01/2018 | 97 |
| X-ray | **5OM1** | 2.1 | Inactive | 1 | CHEMBL202 4114 | Antagonist | 17/01/2018 | 97 |
| X-ray | 6AQF | 2.5 | Inactive | 1 | ZM-241385 | Antagonist | 10/01/2018 | 98 |
| X-ray | **5VRA** | 2.4 | Inactive | 1 | ZM-241385 | Antagonist | 13/12/2017 | 99 |
| X-ray | 5NM2 | 2.0 | Inactive | 1 | ZM-241385 | Antagonist | 27/09/2017 | 100 |

| Method | PDB | Res. (Å) | State | Degree active (%) | Name | Function | PDB Date | Ref. |
|---|---|---|---|---|---|---|---|---|
| X-ray | **8CU6** | 2.8 | Inactive | 1 | LJ-4517 | Antagonist | 31/08/2022 | [82] |
| X-ray | 8CU7 | 2.1 | Inactive | 1 | LJ-4517 | Antagonist | 31/08/2022 | [82] |
| X-ray | **8DU3** | 2.5 | Inactive | 1 | 21a | Antagonist | 10/08/2022 | [83] |
| X-ray | **5NM4** | 1.7 | Inactive | 1 | ZM-241385 | Antagonist | 27/09/2017 | [100] |
| X-ray | 5NLX | 2.1 | Inactive | 1 | ZM-241385 | Antagonist | 27/09/2017 | [100] |
| X-ray | **5MZJ** | 2.0 | Inactive | 1 | theophylline | Antagonist | 26/07/2017 | [101] |
| X-ray | 5MZP | 2.1 | Inactive | 1 | caffeine | Antagonist | 26/07/2017 | [101] |
| X-ray | **5N2R** | 2.8 | Inactive | 1 | PSB36 | Antagonist | 26/07/2017 | [101] |
| X-ray | 5JTB | 2.8 | Inactive | 1 | ZM-241385 | Antagonist | 31/05/2017 | [102] |
| X-ray | **5UVI** | 3.2 | Inactive | 1 | ZM-241385 | Antagonist | 24/05/2017 | [103] |
| X-ray | 5UIG | 3.5 | Inactive | 16 | 5-Amino-N-[(2-Methoxyphenyl)methyl]-2-(3-Methylphenyl)-2h-1,2,3-Triazole-4-Carboximidamide | Antagonist | 08/02/2017 | [104] |
| X-ray | **5K2C** | 1.9 | Inactive | 1 | ZM-241385 | Antagonist | 21/09/2016 | [105] |
| X-ray | 5K2A | 2.5 | Inactive | 1 | ZM-241385 | Antagonist | 21/09/2016 | [105] |

| Method | PDB | Res. (Å) | State | Degree active (%) | Name | Function | PDB Date | Ref. |
|---|---|---|---|---|---|---|---|---|
| X-ray | **8CU6** | 2.8 | Inactive | 1 | LJ-4517 | Antagonist | 31/08/2022 | [82] |
| X-ray | 8CU7 | 2.1 | Inactive | 1 | LJ-4517 | Antagonist | 31/08/2022 | [82] |
| X-ray | **8DU3** | 2.5 | Inactive | 1 | 21a | Antagonist | 10/08/2022 | [83] |
| X-ray | **5K2B** | 2.5 | Inactive | 1 | ZM-241385 | Antagonist | 21/09/2016 | [105] |
| X-ray | 5K2D | 1.9 | Inactive | 1 | ZM-241385 | Antagonist | 21/09/2016 | [105] |
| X-ray | **5G53** | 3.4 | Active | 100 | NECA | Agonist | 03/08/2016 | [50] |
| X-ray | 5IUA | 2.2 | Inactive | 1 | 2-(Furan-2-yl)-5-N-[3-(4-phenylpiperazin-1-yl)propyl]-1H-[1,2,4]triazolo[1,5-a][1,3,5]triazin-8-ium-5,7-diamine | Antagonist | 29/06/2016 | [106] |
| X-ray | **5IU7** | 1.9 | Inactive | 1 | 2-(Furan-2-yl)-5-N-[2-(4-phenylpiperidin-1-yl)ethyl]-1H-[1,2,4]triazolo[1,5-a][1,3,5]triazin-8-ium-5,7-diamine | Antagonist | 29/06/2016 | [106] |
| X-ray | 5IU8 | 2.0 | Inactive | 1 | CHEMBL3934661 | Antagonist | 29/06/2016 | [106] |
| X-ray | **5IU4** | 1.7 | Inactive | 1 | ZM-241385 | Antagonist | 29/06/2016 | [106] |
| X-ray | 5IUB | 2.1 | Inactive | 1 | CHEMBL184061 | Antagonist | 29/06/2016 | [106] |

| Method | PDB | Res. (Å) | State | Degree active (%) | Name | Function | PDB Date | Ref. |
|---|---|---|---|---|---|---|---|---|
| X-ray | **8CU6** | 2.8 | Inactive | 1 | LJ-4517 | Antagonist | 31/08/2022 | [82] |
| X-ray | 8CU7 | 2.1 | Inactive | 1 | LJ-4517 | Antagonist | 31/08/2022 | [82] |
| X-ray | **8DU3** | 2.5 | Inactive | 1 | 21a | Antagonist | 10/08/2022 | [83] |
| X-ray | **4UG2** | 2.6 | Active | 73 | CGS 21680 | Agonist | 08/04/2015 | [107] |
| X-ray | 4UHR | 2.6 | Active | 73 | CGS 21680 | Agonist | 08/04/2015 | [107] |
| X-ray | **4EIY** | 1.8 | Inactive | 1 | ZM-241385 | Antagonist | 25/07/2012 | [58] |
| X-ray | 3UZC | 3.3 | Inactive | 1 | CHEMBL2024114 | Antagonist | 21/03/2012 | [108] |
| X-ray | **3UZA** | 3.3 | Inactive | 1 | compound 4g [PMID: 22220592] | Antagonist | 21/03/2012 | [108] |
| X-ray | 3VG9 | 2.7 | Inactive | 1 | ZM-241385 | Antagonist | 01/02/2012 | [109] |
| X-ray | **3VGA** | 3.1 | Inactive | 1 | ZM-241385 | Antagonist | 01/02/2012 | [109] |
| X-ray | 3PWH | 3.3 | Inactive | 1 | ZM-241385 | Antagonist | 07/09/2011 | [110] |
| X-ray | **3REY** | 3.3 | Inactive | 1 | xanthine amine congener | Antagonist | 07/09/2011 | [110] |
| X-ray | 3RFM | 3.6 | Inactive | 1 | caffeine | Antagonist | 07/09/2011 | [110] |
| X-ray | **2YDO** | 3.0 | Active | 73 | adenosine | Agonist | 18/05/2011 | [53] |

| Method | PDB | Res. (Å) | State | Degree active (%) | Name | Function | PDB Date | Ref. |
|---|---|---|---|---|---|---|---|---|
| X-ray | **8CU6** | 2.8 | Inactive | 1 | LJ-4517 | Antagonist | 31/08/2022 | [82] |
| X-ray | 8CU7 | 2.1 | Inactive | 1 | LJ-4517 | Antagonist | 31/08/2022 | [82] |
| X-ray | **8DU3** | 2.5 | Inactive | 1 | 21a | Antagonist | 10/08/2022 | [83] |
| X-ray | 2YDV | 2.6 | Active | 73 | NECA | Agonist | 18/05/2011 | [53] |
| X-ray | **3QAK** | 2.7 | Active | 73 | UK-432,097 | Agonist | 09/03/2011 | [111] |
| X-ray | **3EML** | 2.6 | Inactive | 1 | ZM-241385 | Antagonist | 14/10/2008 | [112] |

*Table SI 2: Selected performance evaluation metrics generated with 5-fold cross-validation. CV types are StratifiedGroupKFold (PLIF) and StratifiedKFold (ECFP4).*

**PLIF Based RF Classifier Performance Evaluation Metrics**

| #CV | test_roc_auc | test_accuracy | test_precision | test_recall | test_f1 |
|---|---|---|---|---|---|
| 1 | 0.98 | 0.96 | 0.97 | 0.82 | 0.89 |
| 2 | 0.98 | 0.95 | 0.97 | 0.76 | 0.85 |
| 3 | 0.99 | 0.97 | 0.94 | 0.88 | 0.90 |
| 4 | 0.99 | 0.96 | 0.97 | 0.83 | 0.90 |
| 5 | 0.97 | 0.95 | 0.94 | 0.79 | 0.86 |
| **Average** | 0.98±0.01 | 0.96±0.01 | 0.96 ± 0.01 | 0.82±0.02 | 0.88±0.01 |

**ECFP4 Based RF Classifier Performance Evaluation Metrics**

| #CV | test_roc_auc | test_accuracy | test_precision | test_recall | test_f1 |
|---|---|---|---|---|---|
| 1 | 1.0 | 1.0 | 1.0 | 1.0 | 1.0 |
| 2 | 1.0 | 1.0 | 1.0 | 1.0 | 1.0 |
| 3 | 1.0 | 1.0 | 1.0 | 1.0 | 1.0 |
| 4 | 0.98 | 0.97 | 1.00 | 0.88 | 0.94 |
| 5 | 0.97 | 0.98 | 1.00 | 0.93 | 0.96 |
| **Average** | 0.99±0.01 | 0.99±0.01 | 1±0 | 0.96±0.03 | 0.98±0.01 |

*Table SI 3: Predicted probability for the 12 selected compounds to be antagonists according to ECFP4 and PLIF based random forest classifiers.*

| ID | ECFP4-RF Antagonist Probability | PLIF-RF Antagonist Probability |
|---|---|---|
| JG-001 | 0.92 | 0.82 |
| JG-002 | 1.00 | 0.82 |
| JG-003 | 1.00 | 0.82 |
| JG-004 | 0.90 | 0.82 |
| JG-005 | 0.94 | 0.82 |
| JG-006 | 0.99 | 0.82 |
| JG-007 | 1.00 | 0.82 |
| JG-008 | 1.00 | 0.82 |
| JG-009 | 1.00 | 0.82 |
| JG-010 | 0.98 | 0.82 |
| JG-011 | 0.96 | 0.82 |
| JG-012 | 1.00 | 0.82 |

*Table SI 4: Results obtained from our SB approach applied to the best binding poses of the 12 most promising molecules from the virtual screening towards the A2A receptor.*

*values obtained with Kdeep trained against the PDBbind v.2016(REF)

| Ligand Alias | pKd* | Ki (μM) | IC50 (μM) | pIC50 |
|---|---|---|---|---|
| JG-01 | 6.5 | 4.0 | 0.1 | 7.1 |
| JG-02 | 6.2 | 6.6 | 0.1 | 6.9 |
| JG-03 | 5.1 | 94.2 | 1.8 | 5.7 |
| JG-04 | 6.0 | 11.5 | 0.2 | 6.6 |
| JG-05 | 6.0 | 11.3 | 0.2 | 6.7 |
| JG-06 | 5.5 | 35.0 | 0.7 | 6.2 |
| JG-07 | 5.8 | 16.8 | 0.3 | 6.5 |
| JG-08 | 6.0 | 11.3 | 0.2 | 6.7 |
| JG-09 | 5.7 | 24.8 | 0.5 | 6.3 |
| JG-10 | 6.8 | 1.7 | 0.0 | 7.5 |
| JG-11 | 5.7 | 25.2 | 0.5 | 6.3 |

*Table SI 5: Induced Fit redocking of compounds selected by virtual screening workflow.*

| Title | IFDScore XP | GScore |
|---|---|---|
| JG-01 | -584.37 | -13.119 |
| JG-02 | -582.49 | -12.961 |
| JG-03 | -579 | -11.454 |
| JG-04 | -580.23 | -12.589 |
| JG-05 | -582.37 | -12.576 |
| JG-06 | -585.58 | -12.314 |
| JG-07 | -581.77 | -11.601 |
| JG-08 | -582.34 | -14.172 |
| JG-09 | -588.98 | -11.241 |
| JG-10 | -585.45 | -14.55 |
| JG-11 | -587.62 | -13.468 |
| JG-12 | -584.22 | -14.474 |

*Table SI 6: Summary of polar and non-polar contacts of JG-10 in the induced-fit pose.*

Hydrogen Bonds

| Index | Residue | Distance H-A | Distance D-A | Donor Angle | Donor Atom | Acceptor Atom |
|---|---|---|---|---|---|---|
| 1 | PHE168 | 2.8 | 3.2 | 104.93 | 1246 | 2352 [O2] |
| 2 | GLU169 | 2.18 | 3.17 | 175.61 | 1257 | 2352 [O2] |
| 3 | ASN253 | 1.94 | 2.8 | 144.47 | 1863 | 2336 [N1] |

Pi-Stacking

| Index | Residue | Distance | Angle | Offset | Ligand Atoms |
|---|---|---|---|---|---|
| 1 | PHE168 | 3.57 | 8.04 | 0.74 | 2333, 2334, 2337, 2344, 2345 |

Hydrophobic Interactions

| Index | Residue | Distance | Ligand Atom | Protein Atom |
|---|---|---|---|---|
| 1 | ALA63 | 3.6 | 2340 | 473 |
| 2 | ILE66 | 3.74 | 2356 | 495 |
| 3 | VAL84 | 3.48 | 2341 | 621 |
| 4 | VAL84 | 3.54 | 2327 | 621 |
| 5 | LEU85 | 3.6 | 2328 | 629 |
| 6 | PHE168 | 3.88 | 2332 | 1256 |
| 7 | LEU249 | 3.32 | 2331 | 1829 |
| 8 | LEU267 | 3.96 | 2359 | 1971 |

| | | | | |
|---|---|---|---|---|
| 9 | TYR271 | 3.78 | 2357 | 2011 |
| 10 | TYR271 | 3.57 | 2358 | 2009 |
| 11 | ILE274 | 3.71 | 2343 | 2033 |
| 12 | ILE274 | 3.81 | 2359 | 2035 |

*Table SI 7: Tanimoto Similarity Score of candidates selected for experimental testing comparison to most similar known A2A binders found in ChEMBL database[46].*

| ID | $T_c$ | Molecule | Most-similar in ChEMBL | ID | $T_c$ | Molecule | Most-similar in ChEMBL |
|---|---|---|---|---|---|---|---|
| [1] | 0.35 | 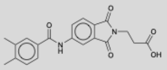 | 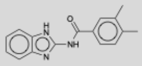 | [7] | 0.67 | 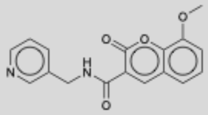 | 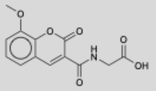 |
| [2] | 0.47 | 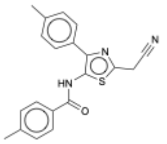 | 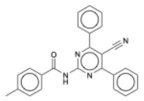 | [8] | 0.31 | 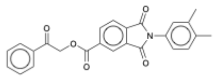 | 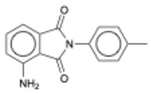 |
| [3] | 0.48 | 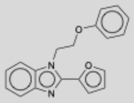 | 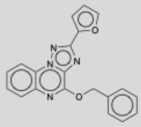 | [9] | 0.35 | 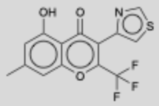 | 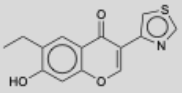 |
| [4] | 0.27 | 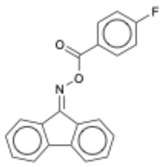 | 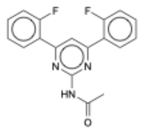 | [10] | 0.33 | 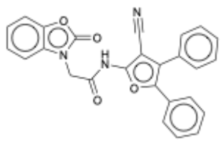 | 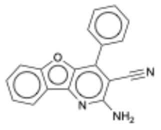 |
| [5] | 0.37 | 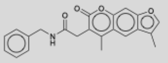 | 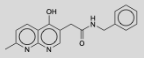 | [11] | 0.33 | 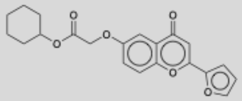 | 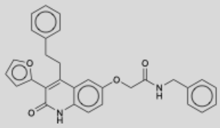 |
| [6] | 0.35 | 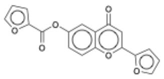 | 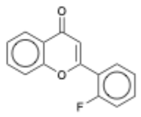 | [12] | 0.36 | 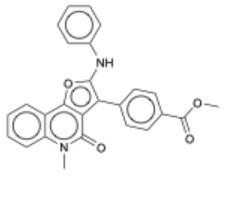 | 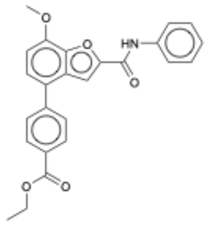 |

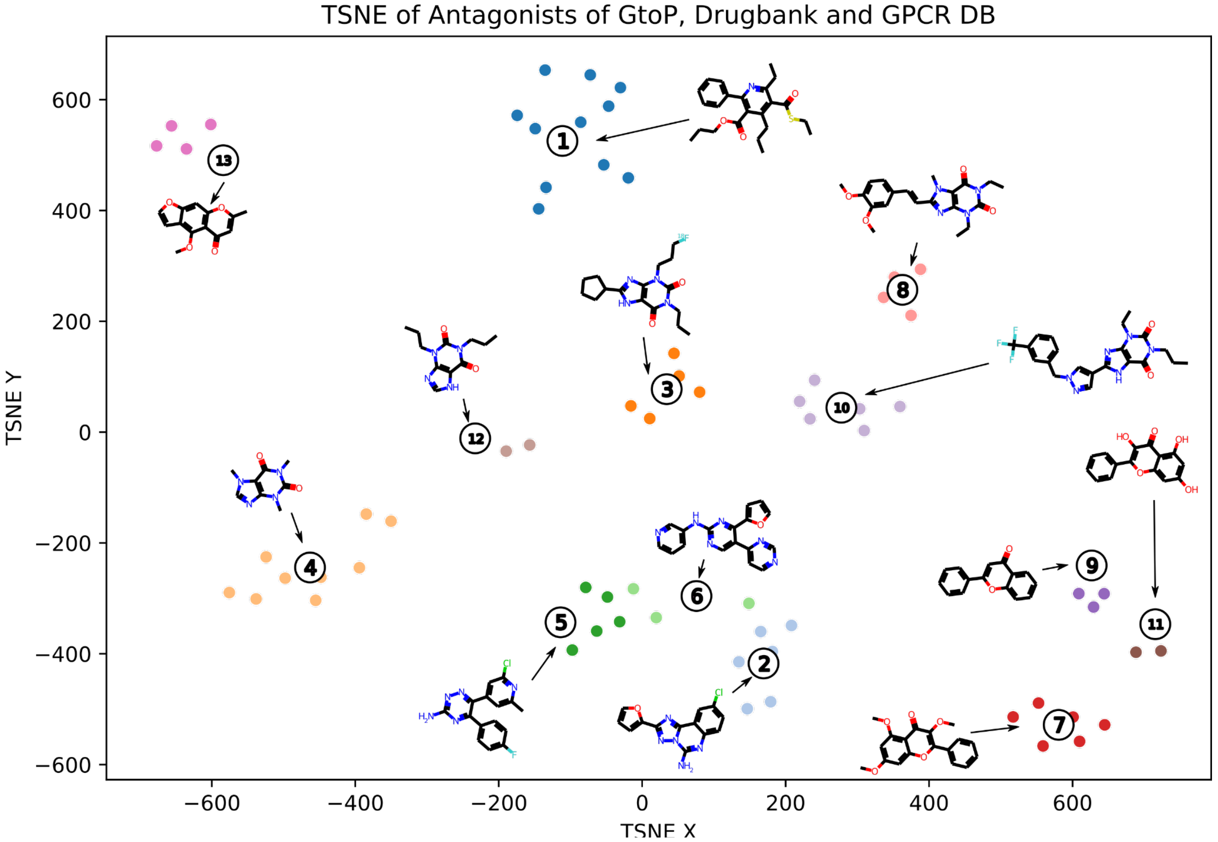


*Figure SI 1: TSNE representation of considered antagonists coming from GtoP, Drugbank and GPCR DB clustered with hierarchical clustering of 13 clusters. Outliers were removed by hand and selected cluster representatives are shown. The chemical space of antagonists is more diverse than that of agonists.*

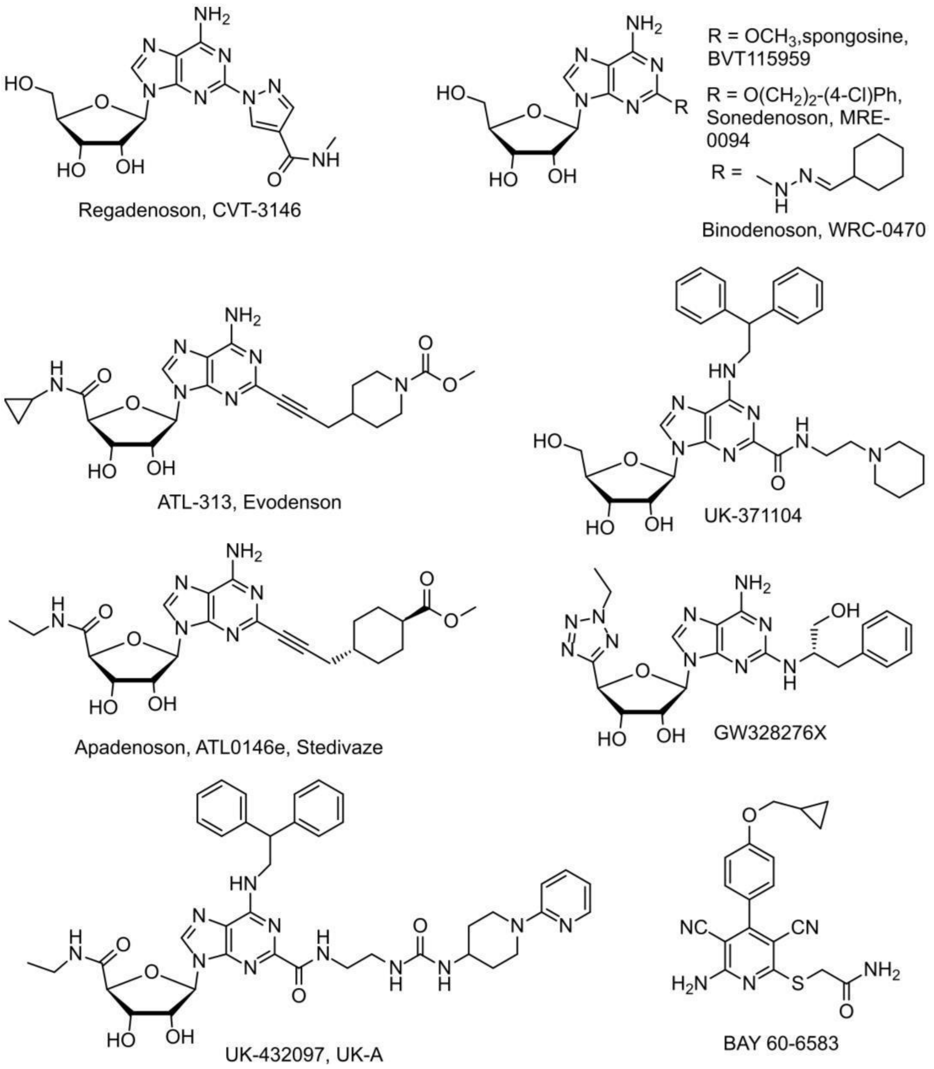


*Figure SI 2: Most of the selective agonists for $A_{2A}$ contain a ribose and an adenine-like moiety.*

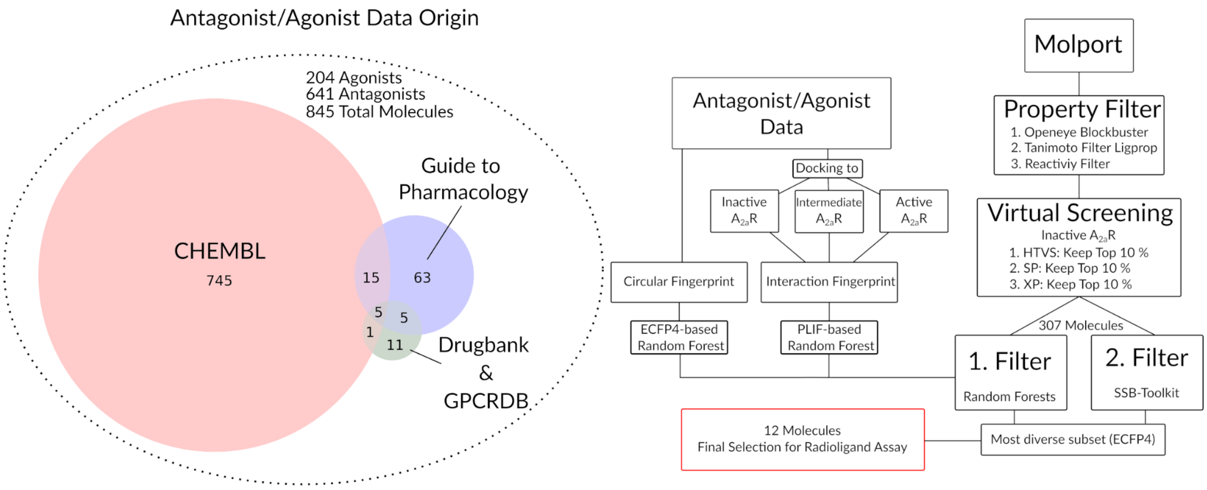


*Figure SI 3: Overview of data collection and overall workflow overview.*

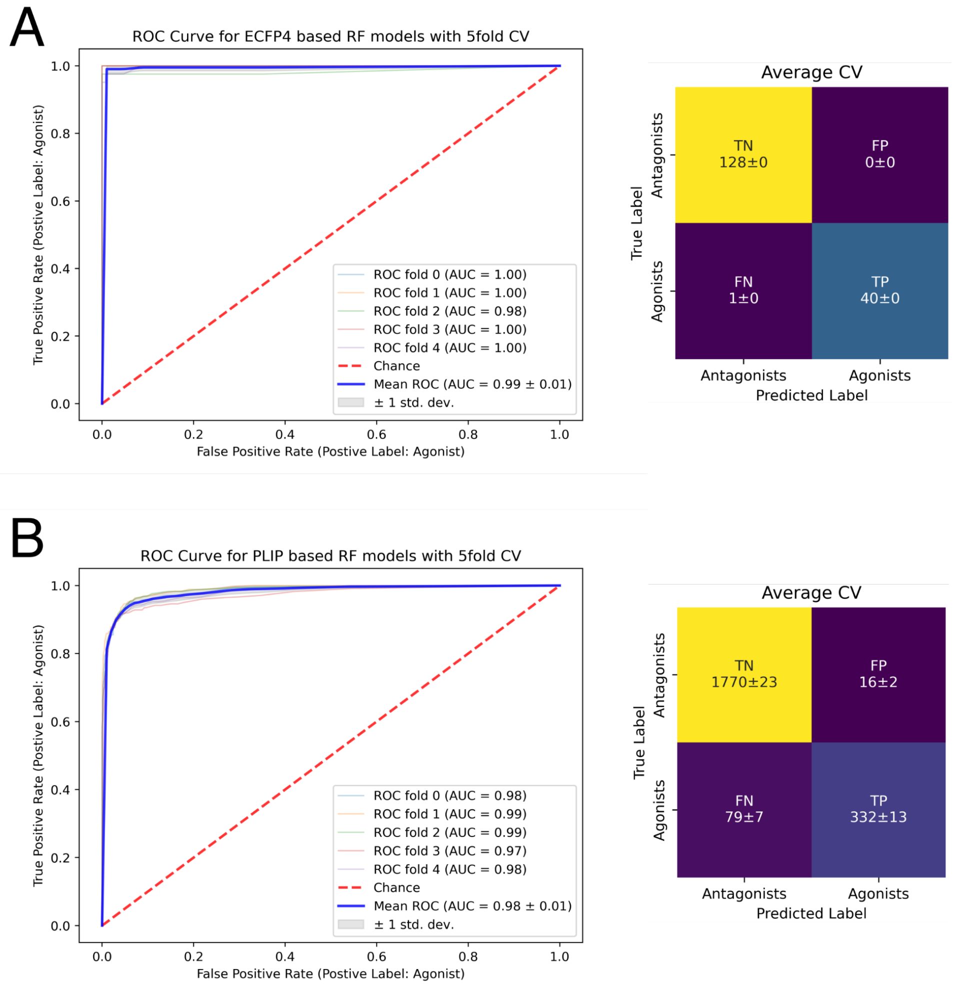


*Figure SI 4: (A, B) Confusion matrix and ROC plot of (A) ECFP4 based and (B) PLIF based random forest model. Validated by 5-fold-cross validation. In the PLIF Based Random Forest model, the best 5 poses for each unique ligand were considered for the classifier. Therefore we applied group cross-validation that ensures that no molecule is shared in between training and test splits. For the ECFP4 model every unique molecule in the training sets was used. The classification by the docking-based approach shows a high accuracy score for the respective splits (compare Table SI 3). Although the area under the curve shows a high value of 0.98, a bit lower than our model based upon the chemical ECFP4 fingerprint, the accuracy of both models is still comparable.*

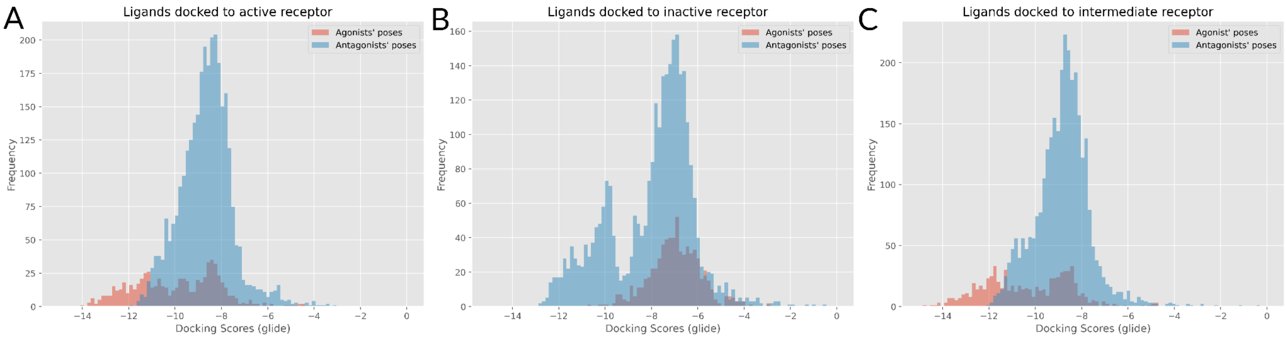


*Figure SI 5: Glide docking scores of the assembled agonist/antagonist library on the three receptors in active(5G53)//inactive(4EIY[58])/intermediate(2YDO) conformation. The average agonist binds with a higher score to the $A_{2A}$ in the active state compared to the inactive state.*

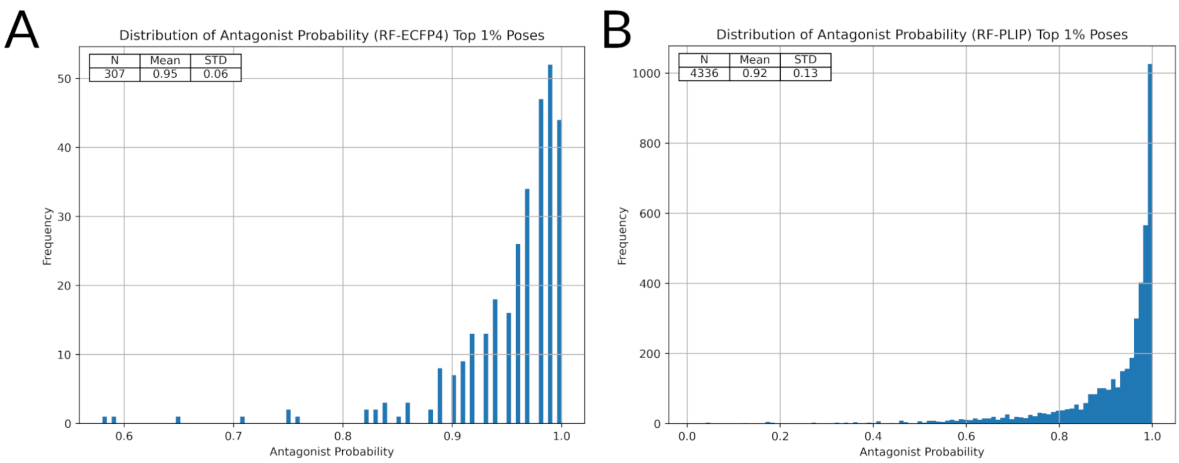


*Figure SI 6: Estimated antagonist probability histograms (100 bins) of top 1% molecules selected by virtual screening to the $A_{2A}R$. A: ECFP4-based RF classifier B: PLIF-based RF classifier. Number of samples (N), arithmetic mean (Mean) and standard deviation (STD) are given for both distributions.*

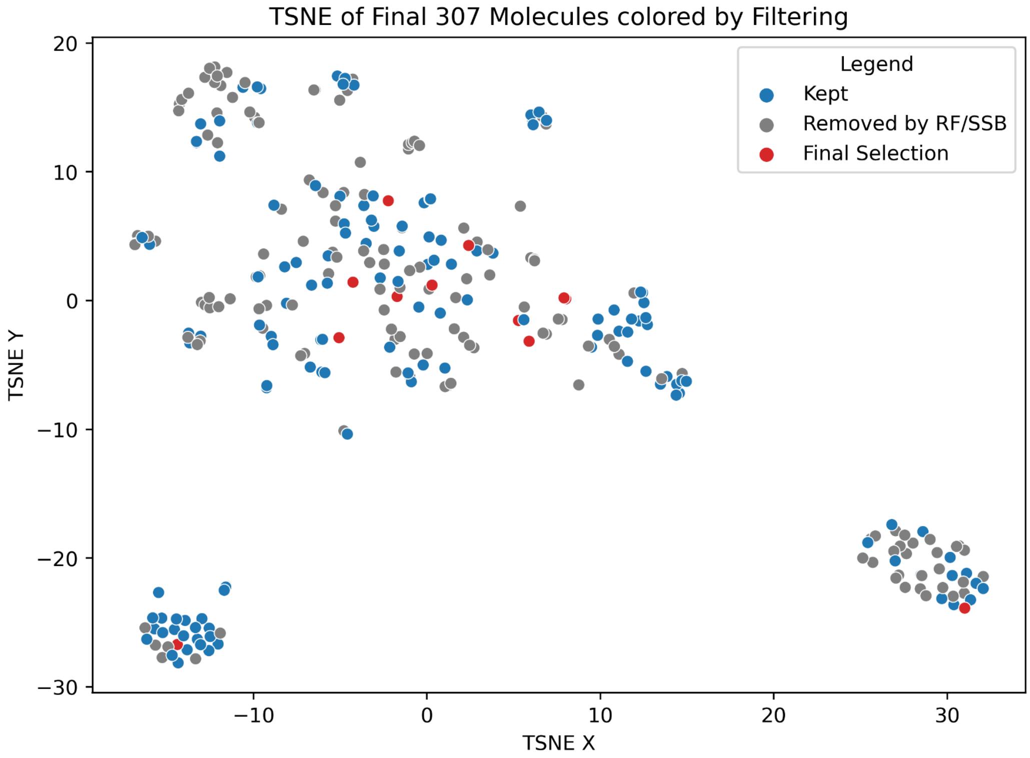


*Figure SI 7: TSNE representation of the 307 molecules identified through virtual screening. Tanimoto similarity was used for the generation of the distance matrix used by the TSNE algorithm. PCA with 50 dimensions was applied.*

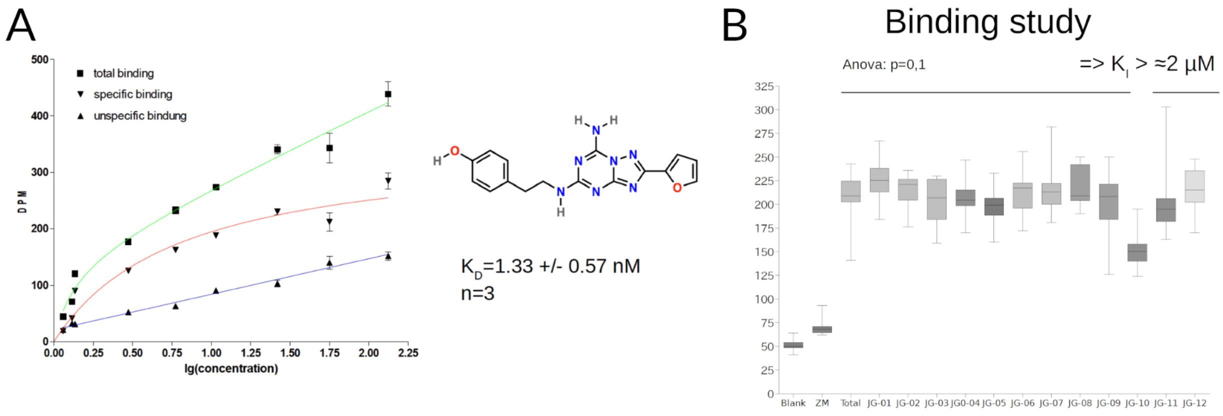


*Figure SI 8: (A) Calibration of assay to gold standard ZMA. (B) Binding study of final 12 selected molecules.*
